# TIST: Transcriptome and Histopathological Image Integrative Analysis for Spatial Transcriptomics

**DOI:** 10.1101/2022.07.23.501220

**Authors:** Yiran Shan, Qian Zhang, Wenbo Guo, Yanhong Wu, Yuxin Miao, Hongyi Xin, Qiuyu Lian, Jin Gu

**Affiliations:** MOE Key Laboratory of Bioinformatics, BNRIST Bioinformatics Division, Department of Automation, Tsinghua University, Beijing, 100084, China; UM-SJTU Joint Institute, Shanghai Jiao Tong University, Shanghai, 200240, China; Department of Automation, Shanghai Jiao Tong University, Shanghai 200240, China

**Keywords:** Spatial transcriptomics, Multimodal information integration, Spatial cluster identification, Gene expression enhancement, Noise reduction

## Abstract

Sequencing-based spatial transcriptomics (ST) is an emerging technique to study in situ gene expression patterns at the whole-genome scale. In addition to transcriptomic data, the technique usually generates matched histopathological images for the same tissue sample. ST data analysis is complicated by severe technical noise; matched histopathological images with high spatial continuity and resolution introduce complementary cellular phenotypical information and provide a chance to mitigate the noise in ST data. Hence, we propose a novel ST data analysis method called transcriptome and histopathological image integrative analysis for spatial transcriptomics (TIST), which integrates the information from sequencing-based ST data and histopathological images. TIST uses a Markov random field (MRF) model to learn the macroscopic cellular features from histopathological images and devises a random-walk-based strategy to integrate the extracted image features, the transcriptomic features and the location information for spatial cluster (SC) identification and gene expression enhancement. We tested TIST both on simulated datasets and on 33 real datasets; we found that TIST achieved superior performance on multiple tasks, which illustrates the utility of this method in facilitating the discovery of biological insights from sequencing-based ST data.

## Introduction

Spatial information is crucial for understanding cell functions in complex physiological and pathological processes. Recent advancements in spatial transcriptomics (ST) technologies have achieved concurrent quantification and localization of mRNA molecules in tissues [1]. Widely used ST techniques can generally be classified into two categories: fluorescence in situ hybridization–based ST (FISH-ST) techniques [2] and sequencing after in situ capture–based ST (SEQ-ST) techniques. SEQ-ST techniques, such as the Visium Spatial Gene Expression platform from 10X, Slide-seq [3] and Seq-Scope [4], are able to interrogate the entire gene set through genome-wide transcriptome profiling.

Despite the obvious advantages of supplementing mRNA sequencing with spatial information, SEQ-ST methods still suffer from technical limitations, including a high mRNA dropout rate and molecular diffusion where mRNA molecules from one spot diffuse to nearby spots during the permeabilization process. These technical factors heavily confound downstream analysis, including feature selection and clustering. Prior studies [5, 6] have shown that joint analysis of ST data and matching histopathological images, generated by haematoxylin and eosin (H&E) staining, has the potential to alleviate the impact of technical factors. Histopathological images complement SEQ-ST in two ways. First, these images have higher resolution and contain embedded cellular phenotypical information. Second, the images are taken prior to ST sequencing; hence, they capture the ground truth cell arrangement without interference from molecular diffusion.

Integrating spatial transcriptomic data with histopathological images remains a challenging problem. Transcriptomic data and histopathological images have different data formats, dimensions, resolutions, and noise levels. It is still an open question as to how to integratively analyse data of both modalities in the identification and classification of spatial clusters (SCs, i.e., targeted tissue regions with distinct gene expression patterns, histological features, and biological functions), as well as in gene expression enhancement, which aims to impute and correct missing and corrupt values of raw ST data affected by under-sampling and molecular diffusion. In this paper, we propose transcriptome and histopathological image integrative analysis for spatial transcriptomics (TIST), an analytical tool suite for ST datasets that includes an SC identification module and a gene enhancement module. Prior works have employed an undirected probabilistic graph model for ST data analytics [7]; TIST expands this model by including network-based data fusion, incorporating transcriptomic, histological and adjacency information into a single spotwise similarity network. From the fused similarity network, TIST identifies and classifies SCs through a customized random walk algorithm and enhances gene expression through a neighbourhood smoothing algorithm. Benchmarks of TIST against state-of-the-art ST analytical methods over simulated and real datasets demonstrate that TIST can drastically reduce the impact of molecular diffusion and high mRNA dropout rates while significantly improving the performance of the expression correlation and spatial differential expression analyses.

## Results

### Overview and evaluation of TIST

Figure 1A illustrates the workflow of the SEQ-ST techniques powered by spatially barcoded probes to capture RNA transcripts in situ and label them with location information. However, due to the limited capture rate and the molecular diffusion that occurs during the permeabilization process, the generated ST data are very noisy. To overcome these drawbacks of the SEQ-ST data, we propose a novel method, designated TIST, to improve the identification of SC and the enhancement of gene expression (Figure 1B) by effectively integrating ST data with histopathological images. By extracting features from the respective modalities, TIST separately builds three modality-specific networks: a histological similarity network generated from high-resolution H&E images through the Markov random field (MRF) technique (Figure S1); a transcriptomic similarity network generated from the gene expression data using the shared nearest neighbour (SNN) method [8]; and an adjacency network converted from spot locations. TIST fuses the three networks into a multi-feature similarity network, termed TIST-net, with SC identification and gene expression enhancement as the main outputs (Materials and methods).

**Figure 1.**
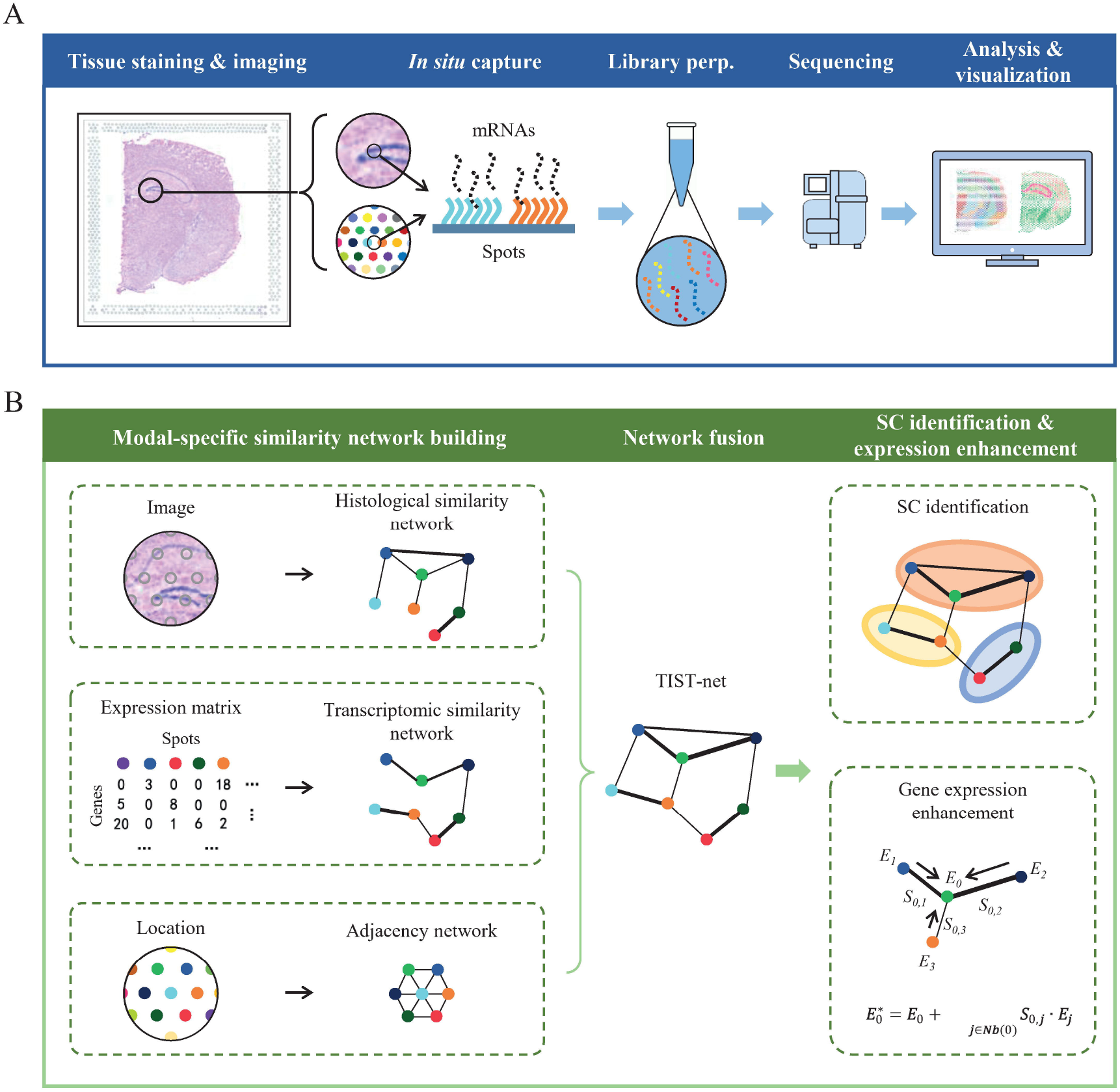
Workflow of SEQ-ST techniques and TIST. **A**. The workflow of popular SEQ-ST techniques. SEQ-ST methods first position and fix tissue sections by printing glass sides with arrayed reverse transcription (RT) primers with unique positional barcodes, with H&E staining and brightfield imaging. Then, mRNA molecules diffuse and hybridize with local RT primers with spatial barcodes. Finally, the spatially barcoded cDNAs were extracted and sequenced for downstream analysis. **B**. The operations in each step of TIST. TIST first obtains multimodal data on histopathological images, RNA sequencing data and location information as input and then extracts histological features, gene expression features, and spot neighbouring information to construct three networks individually, followed by fusing them into TIST-net. Principal outputs of SC identification and gene expression enhancement could be realized through this multi-feature similarity network.

To fully demonstrate the superior performance of TIST on noisy SEQ-ST data, we built noise simulation models (Materials and methods) to systematically test the resistance of TIST to two major types of ST noise: diffusion noise naturally induced during the tissue permeabilization process (Figure 1A) and dropout noise caused by the low capture rate. We applied TIST to a total of 33 real datasets (Table S1). TIST-identified SCs agree well with tissue structures manually labelled with domain knowledge, and TIST-enhanced gene expression profiles could better distinguish the spatial expression patterns than raw data, demonstrating the effective integration of ST data and histopathological images. Overall, our results illustrate the utility of TIST in facilitating the discovery of biological insights from ST data.

### TIST enables unsupervised identification of biologically meaningful tissue domains

Different regions in complex tissues or organs display distinct transcriptomic and histopathological patterns. We applied TIST to two datasets with well-annotated regions and confirmed that TIST could effectively incorporate histological information, leading to biologically meaningful SCs.

We first analysed the mouse cerebral cortex dataset shown in Figure 2A. We used manually annotated tissue domains guided by the Allen Brain Institute reference atlas diagram of the mouse cortex [9] as the ground truth (Figure 2B and Table S2). The structure of the cerebral cortex is highly complex, such that adjacent areas within the cortex might be entirely different in both cell composition and function. For example, the dentate gyrus and the hippocampus proper (Ammon’s horn) are the two main, interlocking parts of the hippocampal formation [10]. The dentate gyrus, consisting of granule cells, contributes to the formation of new episodic memories and is one of the few structures where neurogenesis takes place in the adult brain [11]. The hippocampus proper plays a vital role in learning and memory. The principal cell layers of the two parts are indicated with arrows in the manual annotation results (Figure 2B). We compared the SC identification and partition results of TIST with Louvain [12], a classical clustering method developed for single-cell RNA sequencing (scRNA-seq) data. Figure 2C and D visualize the SC identification results of Louvain and TIST, respectively, over recorded spatial locations (left) and uniform manifold approximation and projection (UMAP) plots (right). On the whole, the processing results of TIST are more consistent with manual annotations than those of Louvain. Specifically, regarding the SC identification of Louvain, the principal cell layer of the dentate gyrus and part of the principal cell layer of the hippocampus proper are incorrectly merged into other parts of the hippocampal formation. Tuning the Louvain parameter does not help improve performance either but leads to fewer contiguous SCs and many more scattered dots (Figure S2). In contrast, TIST agrees well with manual annotation in Figure 2B and achieves good fidelity in more refined tissue structures. The predominant cell layers of the hippocampus and the dentate gyrus were identified and classified into separate SCs, denoted SC_3 and SC_9, respectively, as shown in Figure 2D (left). The 2D UMAP plot in Figure 2D (right) shows that SC_3 and SC_9 are quite similar to the major hippocampal region (SC_2) in transcriptomics, which might explain the failure of Louvain.

**Figure 2.**
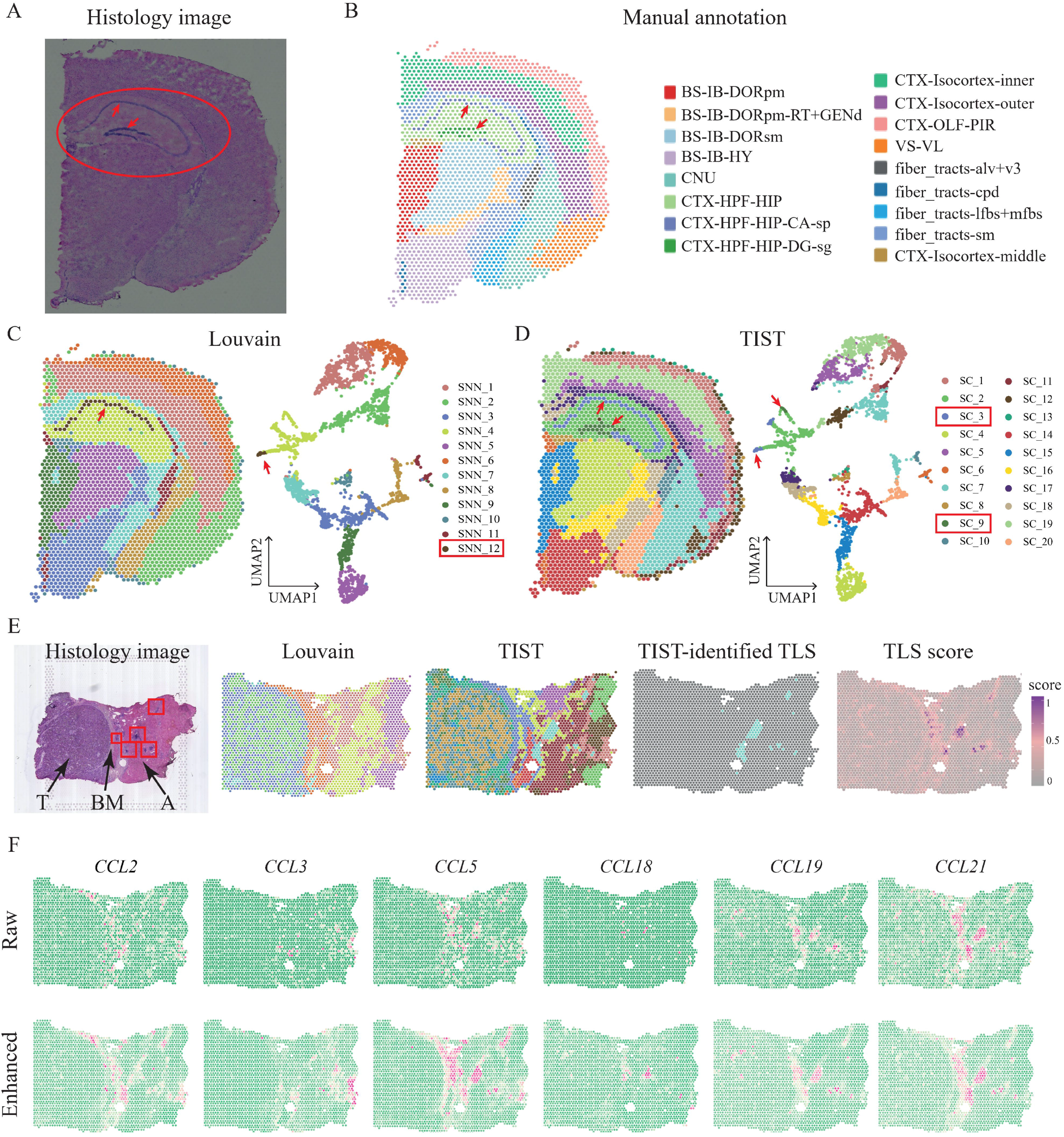
SC identification results of TIST on mouse cerebral cortex dataset and HCC data. **A**. Histopathological image of the mouse cerebral cortex dataset, highlighting the hippocampus proper and the dentate gyrus in the hippocampus formation with red arrows. **B**. Manual annotation of functional domains composing the mouse cerebral cortex as the ground truth. For details on the abbreviations, see Table S2. **C**. SC identification results of Louvain displayed over the spatial locations (left) and the UMAP-transformed 2D pane (right). **D**. SC identification results of TIST displayed over the spatial locations (left) and the UMAP-transformed 2D pane (right). **E**. SC identification results on an HCC dataset. The first column shows the histopathological image of the HCC section, composed of the tumour (T) region on the left, the basement membrane (BM) region in the middle, and the adjacent (A) liver region on the right. Tertiary lymphatic structure (TLS) regions with specific histopathological characteristics are marked with red boxes. The second and third columns show the SC identification results of Louvain and TIST, respectively. The fifth column displays one SC that TIST automatically identifies, which highly matches the TLS regions. The last column visualizes the TLS score of each spot based on the TLS-12 signature scoring method. **F**. Raw expression patterns (upper panel) of typical TLS-12 signature genes, including *CCL2, CCL3, CCL5, CCL18, CCL19* and *CCL21*, and their TIST-enhanced expression patterns (bottom panel).

We further demonstrated the superior performance of TIST with a hepatocellular carcinoma dataset (HCC-1L in Table S1) [13]. The section covers the tumour border, enabling the dissection of immune infiltration. With histological knowledge, we separated the whole section into the tumour region on the left side, the basement membrane in the middle, and an adjacent liver region to the right of the section (Figure 2E). Notably, there are several tertiary lymphatic structure (TLS) regions in the adjacent liver region, marked with red boxes. TLS consists of ectopic aggregates of lymphoid cells in inflamed, infected, or tumoral tissues [14] and are reported to be closely related to tumour progression and metastasis, serving as a potential predictive marker for immunotherapy and prognosis [15]. Accurate identification of TLS regions from ST data is crucial for comprehensively depicting the tumour immune microenvironment. We annotated TLS regions with a TLS-12 signature scoring [16] method as a reference and compared the TLS identification performance of TIST with that of Louvain. As Figure 2E shows, TIST manages to identify a cluster of spatial domains denoted in seafoam colour, which are consistent with annotated high-TLS-score domains based on TLS-12 signature genes. However, the partition given by Louvain has difficulty matching the TLS annotations.

Moreover, we enhanced gene expression patterns through neighbourhood smoothing (Materials and methods) based on TIST-net to interrogate whether the enhanced gene expression could mitigate the technical noise and better recapture the details of biological tissue. Figure 2F shows six genes [16] with reported relationships to TLS and compares their raw expression (upper panel) and enhanced expression (bottom panel). Clearly, the enhanced expressions show much more obvious spatial patterns and match better with the TLS annotation in Figure 2E, confirming the capacity of TIST in integrating ST data with the histopathological image.

### TIST resists the noise induced by mRNA molecular diffusion

The diffusion of mRNA molecules during the permeabilization process brings in inevitable noise challenging ST data analysis but has not received enough attention. In the permeabilization step, mRNA molecules can diffuse from their original location into adjacent spots, leading to inaccurate spatial expression patterns. To showcase the severity of mRNA diffusion, we inspected the empty area in the section without tissue coverage, where no unique molecular identifiers (UMIs) should be present. We observed 1.67%-23.6% (average 8.30%) UMIs detected in empty areas in all datasets (Table S1) and visualized the UMI counts in the empty area of the mouse cerebral cortex dataset in Figure 3A.

**Figure 3.**
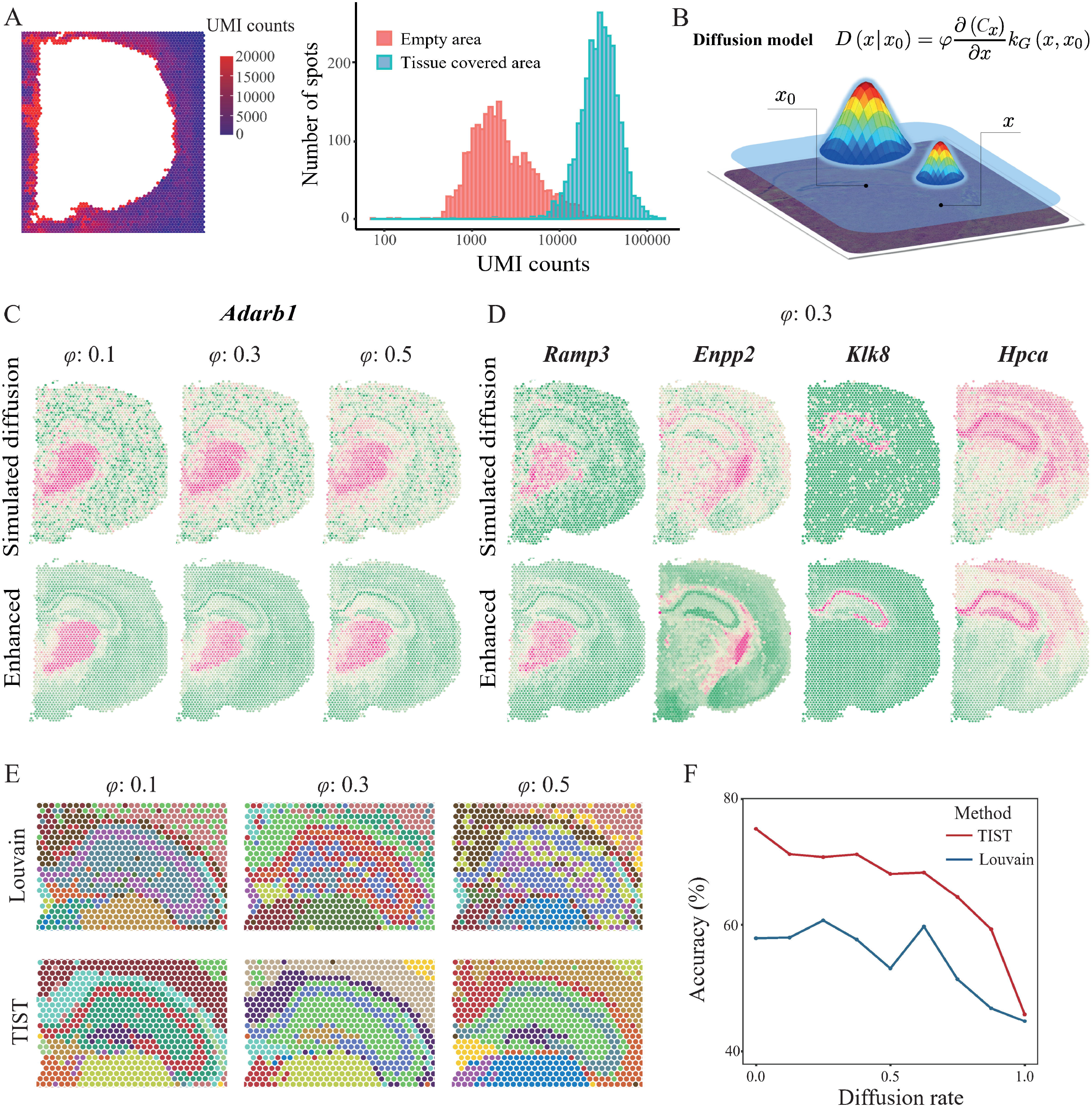
Resistance of TIST to diffusion noise. **A**. Example of mRNA diffusion in SEQ-ST data. The left heatmap displays the UMI counts detected on the slide area not covered by tissue, induced by mRNA diffusion from the tissue-covered area. The right histogram shows the distributions of UMI counts detected in the empty area and in the tissue-covered area on the slide. **B**. Schematic of the simulation model of diffusion noise, describing the diffusion effect that is produced by source spot *x*_*0*_ and acts on any other target spot *x*. The peak height represents the concentrations of mRNA molecules of a certain gene at the matching spot. **C**. Simulated expression patterns of the marker gene *Adarb1* with different settings of diffusion rate (upper panel) and the corresponding TIST-enhanced expression patterns (lower panel). **D**. Simulated expression patterns of the marker genes *Ramp3, Enpp2, Klk8*, and *Hpca*, with a diffusion rate fixed at 0.3 (upper panel) and the corresponding TIST-enhanced expression patterns (lower panel). **E**. Performance of Louvain (upper panel) and TIST (lower panel) in identifying and segregating the hippocampus proper and the dentate gyrus at different simulated diffusion rates. **F**. SC identification accuracies of Louvain (blue line) and TIST (red line) on the whole dataset given different simulation diffusion rates.

We developed a diffusion simulation method according to Fick’s first law [17] of diffusion (Materials and methods) and benchmark TIST on simulated datasets with ground truth. Figure 3B shows that the simulated mRNA diffusion effect from source spot *x*_*0*_ to any other target spot *x*, denoted *D* (*x*|*x*_*0*_), is negatively related to the distance between the two spots and positively related to the concentration gradient and diffusion rate *φ*. Here, the diffusion rate *φ* models the influence of potential diffusion-related factors, including temperature, liquid viscosity and molecular size.

We showed resistance of TIST to diffusion noise with marker genes that are specifically expressed in certain tissue domains. *Adarb1*, reported to be associated with the nervous system [18], was found to be specifically expressed in the sensory-motor cortex–related zone of the thalamus (BS-IB-IORsm). We tested simulated *φ* ranging from 0.1 to 0.9 and exhibit the results in Figure 3C and Figure S3A. TIST-enhanced expression of *Adarb1* restored its spatial pattern even when the diffusion effect was tremendous (Figure 3C). We further investigated various genes marking different SCs with fixed diffusion rates (Figure 3D and Figure S3B). Comparing TIST-enhanced profiles with raw expression patterns (Figure S4) not undergoing any simulation diffusion, we could conclude that TIST-enhanced ST data well reserve the spatial expression patterns and enable stable recapitulation of tissue structures even with severe diffusion noise.

We further benchmarked TIST for SC identification and classification with simulated datasets at different diffusion rates and compared TIST with Louvain. The performance of TIST compared with Louvain in identifying the hippocampus proper and the dentate gyrus is shown in Figure 3E and Figure S3C. To fully evaluate SC identification performance of resisting diffusion noise, we investigated whether TIST could identify SCs from simulated diffusion datasets well resembling manually annotated results in Figure 2B. As Figure 3F shows, TIST indeed achieves superior SC identification and classification performance compared to Louvain with higher recognition accuracy across all diffusion rate settings. An unsupervised modularity index was also adopted as the metric [19] and agreed that TIST performs better than Louvain at SC identification with diffusion noise (Figure S3D).

### TIST accommodates high dropout rates

As with scRNA-seq, ST methods also suffer from a low mRNA capture rate, resulting in a large proportion of false zeros for expressed genes, which are called “dropout” events. Currently, SEQ-ST datasets can reach a median of 0.4k to 6k genes detected per spot (Table S1). A high ratio of dropout would compromise the accuracy of downstream analyses. There is an urgent need to develop ST analysis methods to overcome high dropout rates.

First, *Hpca*, a specific marker gene of the hippocampus proper [20], was selected to test whether TIST could enhance gene expression effectively from ST data with different dropouts. We randomly removed 10%-90% of non-zero data in the original expression matrix and used TIST-net to fill and restore the expression patterns. As the dropout rate increased, the aggregation effect of *Hpca* expression gradually weakened (Figure 4A, upper panel). Its spatial expression pattern became increasingly difficult to discern based on raw data. TIST-enhanced expression maintained distinct domain enrichment until the dropout rate reached 0.8 (Figure 4A, bottom panel and Figure S4). Fixing the simulated dropout rate as 0.5, we further inspected various marker genes; the results are shown in Figure 4B and Figure S5A. In addition, we designed a recovery score to quantitively measure how well TIST-enhanced expression profiles from simulated dropout datasets resemble expression patterns before simulating dropouts (for details, see Materials and methods). We selected 49 genes (Table S3) reported to be related to brain activities [9] for evaluation. As Figure 4C shows, it would be difficult to recover gene expression with high fidelity if we dropped more than 40% of the expressed genes from the currently measured ST data.

**Figure 4.**
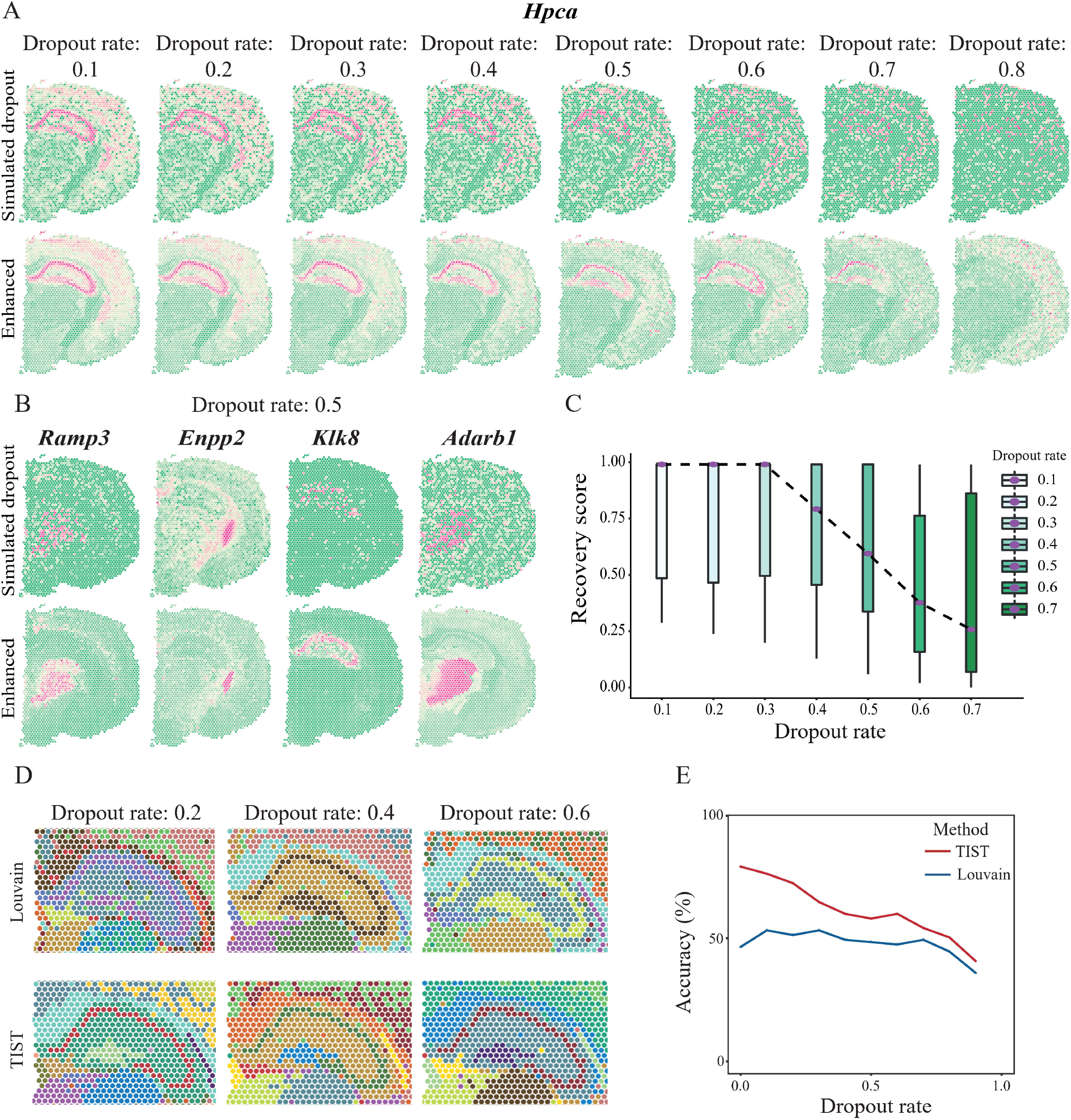
Resistance of TIST to dropout noise. **A**. Simulation expression patterns of the marker gene *Hpca*, with dropout rates ranging from 0.1 to 0.8 (upper panel) and the corresponding TIST-enhanced expression patterns (lower panel). **B**. Simulation expression patterns of the marker genes *Ramp3, Enpp2, Klk8*, and *Adrab1* at a fixed dropout rate of 0.5 (upper panel) and the corresponding TIST-enhanced expression patterns (lower panel). **C**. Boxplot of recovery scores TIST achieves on 49 genes reported to be related to brain activity under different simulation dropout rates. The dotted line connects the median recovery scores in each condition. **D**. Performance of Louvain (upper panel) and TIST (lower panel) in identifying and segregating the hippocampus proper and the dentate gyrus at different simulation dropout rates. **E**. SC identification accuracies of Louvain (blue line) and TIST (red line) on the whole dataset given different simulation dropout rates.

Next, we evaluated the influence of dropout on SC identification. Using the example of the hippocampus proper and the dentate gyrus, we compared the performance of TIST against Louvain with a simulated dropout rate ranging from 0.1 to 0.8. As shown in Figure 4D, Louvain always fails to distinguish the two regions, while TIST allows accurate identification and segregation of the two regions when the simulated dropout rate is less than 0.6. We also quantitatively measured SC identification performance on the whole mouse cerebral cortex dataset with both the supervised and unsupervised metrics (Figure 4E and Figure S5B), finding that the dropout resistance of TIST regarding SC identification significantly surpassed that of Louvain.

### TIST refines the detection of spatial expression patterns

The detection of spatially differentially expressed genes (SDE genes) is a powerful way to link spatial regions with biological functions. Previous work has shown that SPARK, built upon a generalized linear spatial model with a variety of spatial kernels, is a powerful tool for revealing spatial expression patterns [21]. We thus compared TIST with SPARK in identifying SDE genes on the mouse cerebral cortex dataset.

We evaluated SDE gene identification by inspecting the top 100 candidate SDE genes given by TIST and SPARK (Materials and methods). As shown in Figure 5A, 13 genes were commonly included in the top 100 by TIST and SPARK. We visualized the expression patterns of typical genes to assess the quality of the identified SDE genes. Figure 5B shows several instances of commonly identified genes, all displaying clear spatial patterns. *Lypd1*, a marker of sensory areas [22], was successfully identified by both methods. Figure 5C shows typical genes of the 87 genes among the top 100 by TIST only, all well recapitulating the spatial tissue structure. *Hpca*, a marker of the hippocampus proper, was highly ranked by TIST as an SDE gene but was not considered significant by SPARK (p value=0.85). In contrast, 87 genes recognized by SPARK displayed either scattered or singularly high expression patterns, as shown in Figure 5D. We then defined a spatial enrichment score (Materials and methods) to quantitatively evaluate the quality of SDE genes. TIST identified the top 100 candidates with significantly higher spatial enrichment scores than SPARK (Figure 5E), proving the superior performance of TIST at SDE gene identification. We further investigated the functional annotation of the 87 exclusive genes identified by TIST and SPARK and showed the results in Figure S6. Gene set enrichment analysis (GSEA) with Gene Ontology (GO) pathways showed that the 87 SDE genes identified by TIST were significantly enriched in pathways related to brain activity. The 87 SDE genes of SPARK, however, are closely related to the functions and activities of the heart, which is difficult to explain. In summary, TIST could resist technical noise and warrant the detection of spatial expression patterns biologically and functionally related to the tissue.

**Figure 5.**
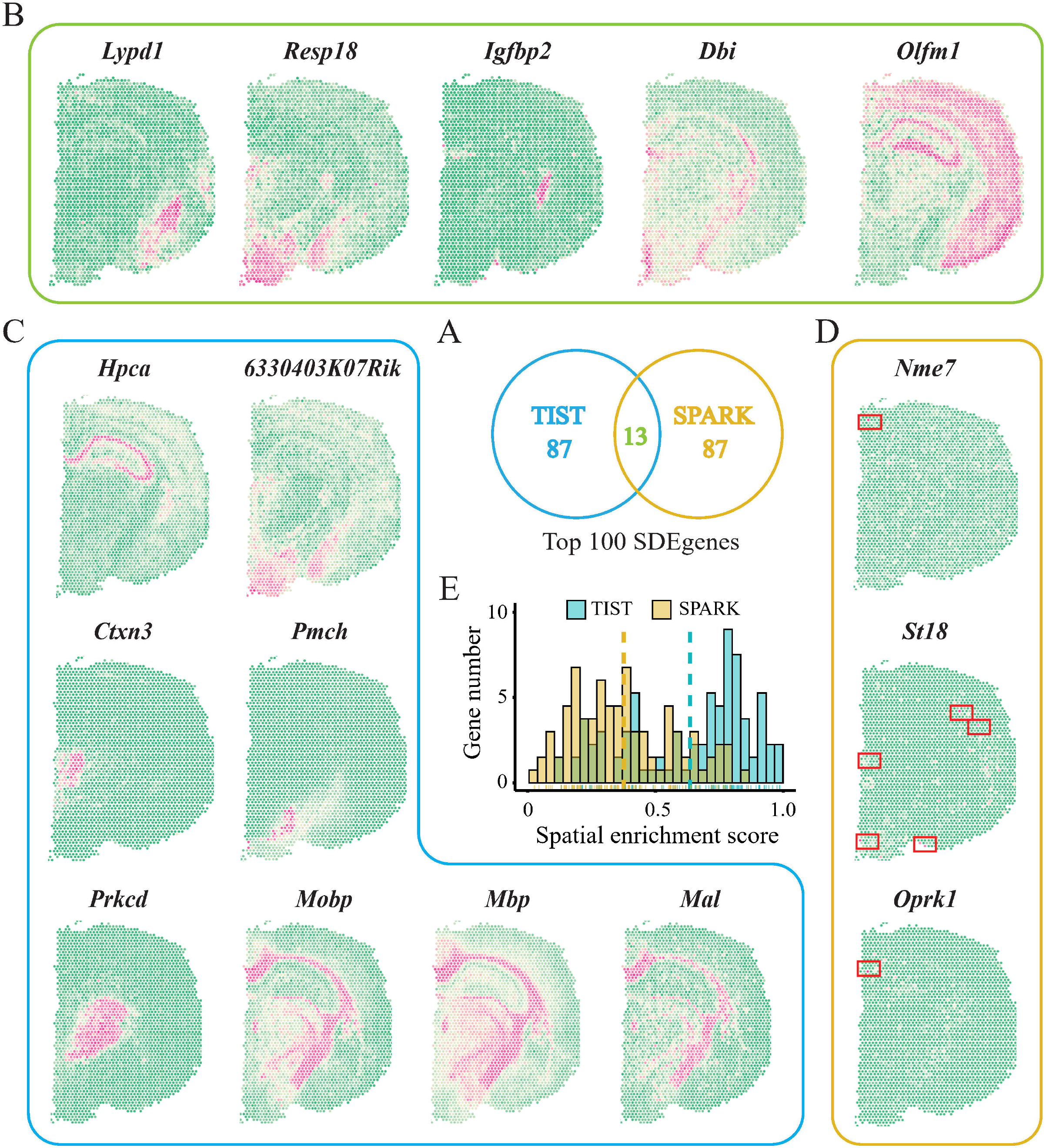
Performance of TIST in facilitating SDE gene detection. **A**. Venn diagram of the top 100 SDE genes identified by TIST and SPARK. The colour of each part corresponds to the border line of **B, C**, and **D. B**. Spatial expression patterns of typical genes commonly identified by both TIST and SPARK in the top 100 SDE genes. **C**. Spatial expression patterns of typical genes detected by TIST, but not by SPARK, in the top 100 SDE genes. **D**. Spatial expression patterns of typical genes detected by SPARK, but not by TIST, in the top 100 SDE genes. Scattered spots with singularly high expressions are marked with red boxes. **E**. Quantitative assessment of the top 100 SDE genes detected by TIST and SPARK.

### TIST benefits gene co-expression inference from ST data

It remains a great challenge to effectively incorporate spatial information from ST data and enable reliable detection of gene co-expression patterns in specific tissue regions. We show that TIST-identified SCs could facilitate the detection of region-specific gene co-expression patterns and shed light on complicated cell–cell communication in tissue.

The hippocampus formation in the mouse cerebral cortex dataset is split by TIST into three SCs, including principal cell layers in the hippocampus proper (SC_3), the dentate gyrus (SC_9) and the remaining part (SC_2). We first compared the gene co-expression strength within the three SCs. Figure 6A visualizes the Spearman correlation values between any pair of genes in the top 2,000 SDE genes found by TIST. Gene co-expression is significantly stronger in the dentate gyrus region (SC_9) than in the other two regions, suggesting that the dentate gyrus undergoes highly precise regulation of gene expression, which might account for its critical role in neurogenesis. We further investigated cell communication with *CellChat* [23] to infer the strength of cell communication within and among the three SCs. Figure 6B shows significant ligand–receptor pairs within each SC, among which we marked cases supported by literature evidence with red underlines (Table S4). For instance, *Wnt* ligands binding to the combination of *FZD1/3* and *LRP6* co-receptors are reported to activate intracellular signalling and facilitate brain development as well as adult hippocampal neurogenesis [24], in line with facts that the dentate gyrus region (SC_9), where neurogenesis takes place, enriches up to four related ligand–receptor pairs (marked with red box). Figure 6C visualizes significant ligand–receptor binding patterns among the three SCs. Specifically, the hippocampus proper (SC_3) communicates quite actively with the other two SCs, as the red colour representing SC_3 takes up the highest proportion of ligand–receptor pairs in the outer ring. We further analysed the communication between the three SCs and the other TIST SCs identified in the mouse cerebral cortex dataset. SC_3 maintained generally strong interactions with most SCs (Figure 6D). These observations echo the fact that the hippocampus proper contains hub neurons possessing widespread axonal arborizations [25].

**Figure 6.**
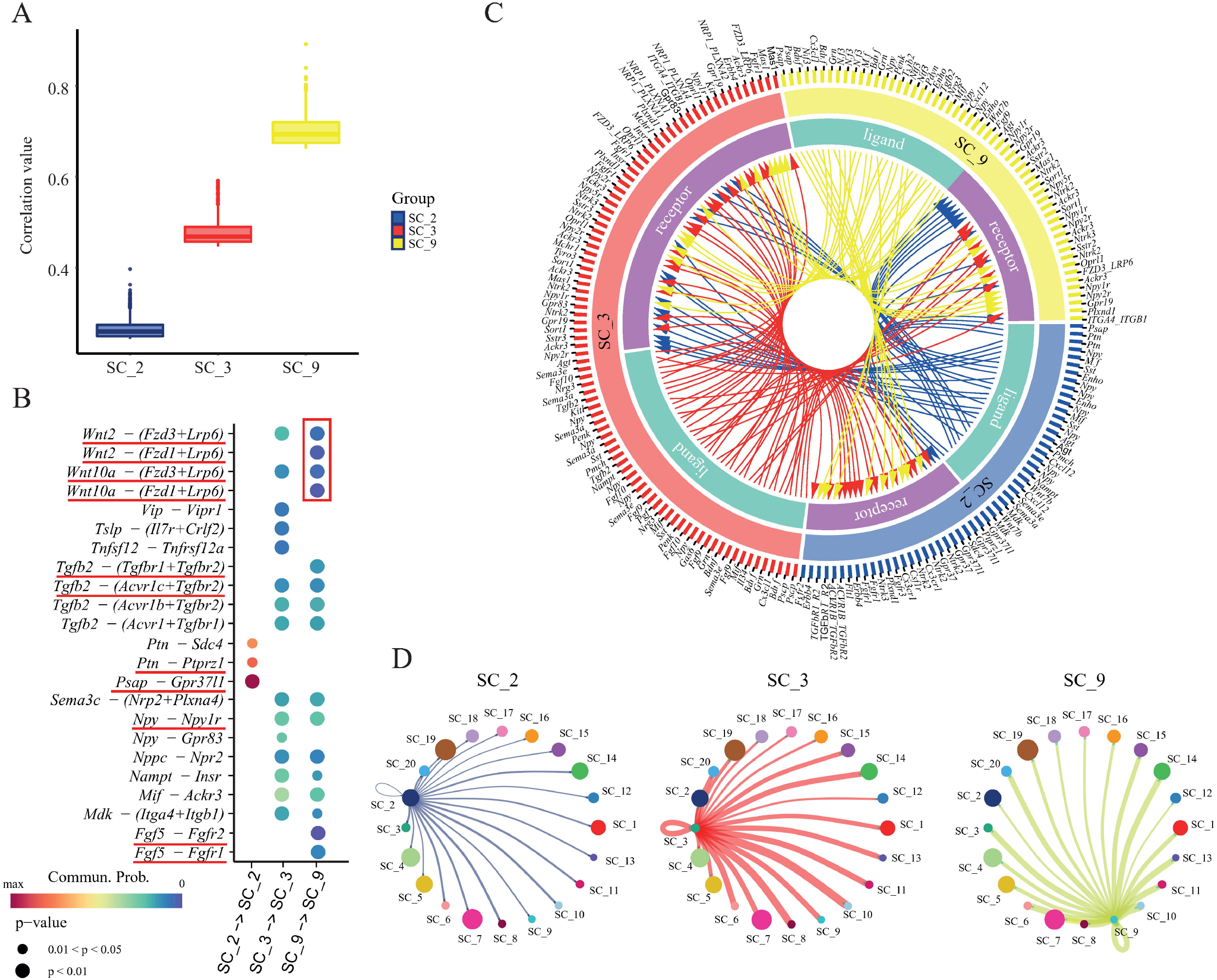
Spatial co-expression analysis based on TIST. **A**. Bar plot showing the co-expression strength (Spearman correlation) within three SCs (SC_2, SC_3 and SC_9) composing the hippocampal formation. **B**. Bubble plot of ligand–receptor pairs enriched within the three SCs. The area of the bubble represents the p value, while the colour shows the interaction strength. We underline the ligand–receptor pairs supported by literature evidence. **C**. Circos plot showing ligand–receptor pairs among SC_2, SC_3 and SC_9. Curved arrows indicate the ligand–receptor relationships. Genes detected to be specific in different SCs are listed outside the circle. **D**. Communication strength between the three SCs and the others. The width of the curve represents the communication strength between two SCs that the curve connects. Spot size represents the number of enriched ligand–receptor pairs.

### TIST adapts to diverse tissue types and spatial resolutions

In addition to the mouse cerebral cortex dataset and the HCC dataset shown in Figure 2, we benchmarked TIST with 33 public ST datasets in total (Table S1), covering diverse tissue types, different pathological conditions and spatial resolutions, demonstrating the versatility of TIST. In detail, there were 12 public 10X ST datasets (Figure 7A), 21 public 10X ST datasets from our previous study [13] (Figure 2E and Figure S7) and 1 Seq-Scope dataset of mouse liver (Figure 7B). The results in Figure 7A show that TIST always managed to recapitulate delicate tissue structures regardless of the tissue type or pathological condition. Figure 7B (left) shows the histopathological image of a normal mouse liver sample and marks the tissue region for ST sequencing by Seq-Scope with a resolution of 10 µm. TIST-identified SCs (right) highly resemble the annotation from the original paper (middle). In summary, TIST is platform-independent and adapts well to diverse sample sources and resolutions.

**Figure 7.**
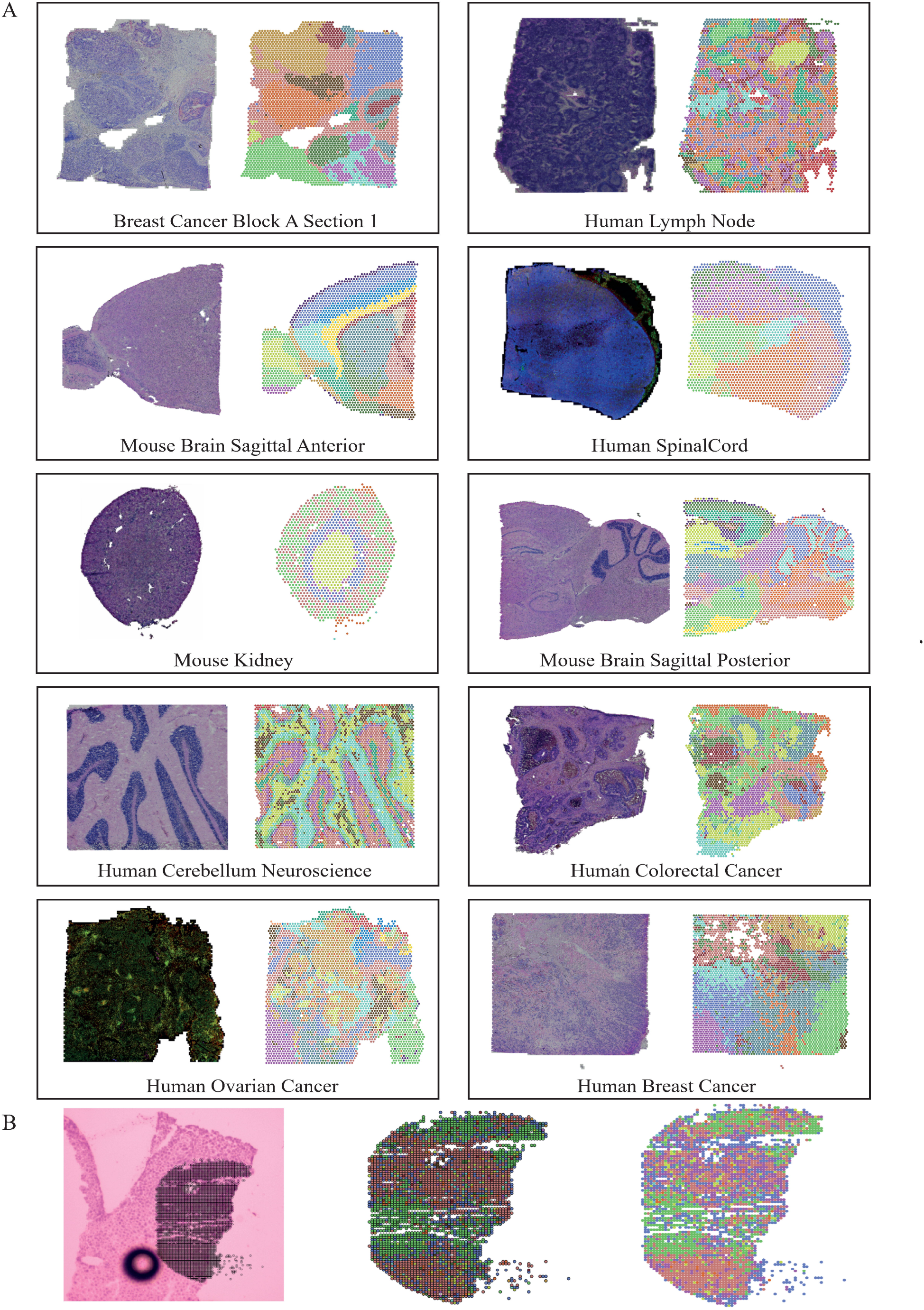
SC identification results of TIST on datasets with diverse sample types and spatial resolutions. **A**. SC identification results of TIST on public datasets from 10X. Each black box corresponds to one dataset. Figures on the left side within the boxes show the histopathological images. The right figures within the boxes show the automatic SC identification results of TIST. **B**. SC identification results of TIST on the mouse normal liver sample sequenced by Seq-Scope. The histopathological image is shown on the left with a dark mark denoting the region for ST sequencing. The middle figure shows the SC identification results given by the original paper. The right figure shows the SC identification results of TIST.

## Discussion

We proposed TIST tailored towards SEQ-ST data by effectively integrating transcriptome and histopathological images via a network-based multimodal data fusion strategy. TIST could identify SCs of spots with distinct gene expression patterns, histological features and biological functions in an unsupervised manner and enhance spatial expression signals from noisy ST data. The design of TIST warrants its adaptability to different spatial resolutions and compatibility with diverse ST techniques once the histopathological image, transcriptome and location information are all available.

We reported that mRNA diffusion during the process of tissue permeabilization inevitably induces noise in ST data and blurs spatial expression patterns, which has not yet received enough attention. We demonstrated the severity of the diffusion noise and its interference in downstream ST analyses and showed that TIST could mitigate the technical noise by effectively integrating ST data and the matching histopathological images.

Joint analysis of ST data and histology is reported to have the potential to alleviate the impact of technical factors. Methods recently proposed to account for the spatial dependency of gene expression and cellular phenotypical information from histopathological images are mainly based on deep learning models, including stLearn [5] and SpaGCN [7]. Instead, we leverage the classic image feature extraction technique, MRF, which has been demonstrated to have good utility in processing H&E images, to extract informative histological features of individual spots. We represent the similarities from histology, transcriptomics and adjacency into three networks and fuse them into a single network via a devised random-walk-based strategy. We benchmarked TIST against stLearn and SpaGCN on a mouse posterior brain dataset (Figure S8) and showed that TIST is able to detect and segregate subtle tissue structures that are missed by stLearn and SpaGCN, suggesting that TIST provides a superior framework to integrate multimodal ST data.

The rapid development of ST techniques has relieved some technical drawbacks, such as limited resolution, but many challenges remain to be solved [26]. Ideally, we want to obtain the expression profiles of individual cells together with their locations. For SEQ-ST, each spot captures mRNAs from proximal cells in the tissue section covering the slide. It would be difficult to obtain the mapping relationship between spots and cells and accurately recover expression profiles at the single-cell level from spot-level ST data. As histopathological images are taken prior to ST sequencing, they are expected to capture the ground truth cell arrangement and help obtain ideal single-cell ST data. It may be possible to extend TIST to realize accurate single-cell ST analysis given that TIST provides an excellent multimodal feature extraction and data fusion method.

## Materials and methods

### Data availability

The R package TIST is an open source available at http://lifeome.net/software/tist/. We used 33 datasets, including 12 10X Genomics published datasets, 21 10X human liver datasets from our previous study, and 1 mouse normal liver sample processed by Seq-Scope in this study. Among these samples, one mouse cerebral cortex sample and one HCC sample were used to assess the performance of TIST in depth. The 12 10X Genomics datasets were downloaded from 10X genomics support (https://www.10xgenomics.com/resources/datasets); the Seq-Scope dataset was obtained from the Gene Expression Omnibus (GEO) under accession number GSE169706, and the human liver datasets were downloaded from GSA under accession number HRA000437.

### Data preprocessing

To extract the effective pixels of H&E staining images, we first converted them into greyscale and then fitted a mixed Gaussian model to identify the foreground pixels covered by tissue. For spatial RNA-seq data. We normalized the gene expression using the R package *Seurat* (4.0.3) by the default method *LogNormalize*. Then, the expression of each gene was scaled to a standard normal distribution.

### TIST: Construction of image-based histological similarity network

We then constructed a network to extract features of histopathological images. First, a Markov random field (MRF) model was used to assign a label to each pixel. Pixels were pre-clustered by *k*-means into 50 clusters as default. The labels were iteratively updated by considering the 3*3 neighbours using MRF with 10 runs.

Since each spot covers multiple pixels, we converted the pixel labels that belonged to a spot as a 50-dimensional probability distribution as the spotwise features. Then, we constructed the image-based spotwise similarity network by calculating the symmetrical Kullback–Leibler (KL) divergences *L*_*u,v*_ between label distributions of any two spots *u* and *v*:

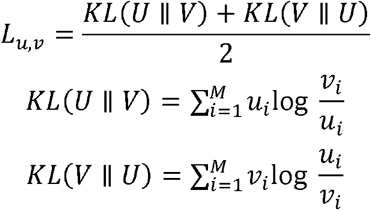

Here, *U* and *V* are the spotwise image feature vectors (1*50, as *u*_*i*_ and *v*_*i*_ represent the probability that each pixel belongs to a certain label *i*), and *M* represents all labels in a spot. Then, the adjacency matrix of the histological similarity network could be calculated as follows:

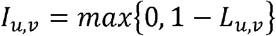

### TIST: Construction of transcriptomic similarity network by SNN

We next constructed a spotwise transcriptomic similarity network. First, the top 2,000 highly variable genes were selected by the *FindVariableGenes* function. Then, we performed principal component analysis (PCA) on the scaled data of the selected genes. Finally, the top 50 principal components (PCs) were used to construct an SNN [8] graph using the *FindNeighbors* function with the default settings. The edge weights of the SNN graph were used as the adjacency matrix of the transcriptomic similarity network:

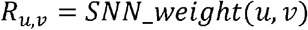

### TIST: Construction and integration of the multi-feature similarity network

Finally, we integrated the above two networks from images and RNA-seq data as a multi-feature network (designated TIST-net). TIST-net used the transcriptomic similarity network to define the network topology (retaining only the edges in the transcriptomic similarity network). Additionally, the physical spatial distances between spots need to be considered (Manhattan distance *DM*_*u,v*_ was used in this study). The adjacency matrix of TIST-net was defined as follows:

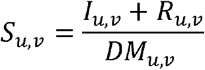

*S*_*u,v*_ was then normalized to 0-1 by:

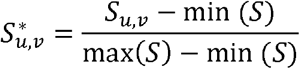

### TIST: Enhancement of gene expression by neighbourhood weighting

Using the neighbouring information in TIST-net, we could enhance gene expression by calculating the weighted average of the expression of immediate neighbours for a single spot. For any spot *u* and *v*, we define:

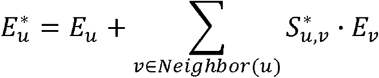

Here, *E*^*\**^ is the enhanced gene expression, and *E* is the raw gene expression. *Neighbor*(*u*) represents the immediate neighbours of spot *u* in TIST-net.

### TIST: Identification of spatial clusters by Walktrap

We used the Walktrap algorithm [27] to identify spatial clusters based on random walks to make full use of neighbouring relationships within networks. Here, we define the transition probability between different states as *p*_*uv*_, the probability of random walk from *i* to *j* is *P*_*uv*_, and calculate the random walk distance *DW*_*uv*_ between spots *u* and *v* as follows:

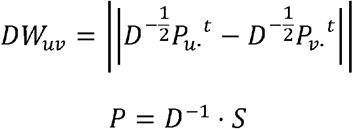

where *D* is the diagonal matrix of the degrees; *t* represents the step of the random walk; and *S* is the edge weight of TIST-net.

### Diffusion effect simulation experiments

To mimic the diffusion effect of the permeabilization process, we used *D* (*x*|*x*_*0*_) to denote the diffusion effect produced by source spot *x*_*0*_ and acting on any other target spot *x*, based on Fick’s first law of diffusion [17]. We assume that each spot diffuses to other spots with lower concentrations and define:

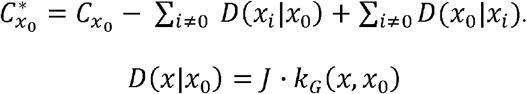

where *C*_*xo*_ represents origin gene counts in spot *x*_*0*_; *C*^*^_*XO*_ represents gene counts in spot *x*_*0*_ after diffusion, and *D* (*x*|*x*_*0*_) is the gene counts in diffusion from spot *x*_*0*_ to spot *x*. Based on Fick’s first law, *J* represents diffusion flux, defined as:

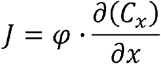

where *φ* represents the diffusion rate, which is related to temperature, liquid viscosity, and molecular size. We adjusted *φ* in the range {0.1, 0.2, 0.3…, 0.9} to obtain different simulation results. *k*_*G*_ (*x*, *x*_*0*_) is the Gaussian kernel function on the distance matrix, which can be written in the following form:

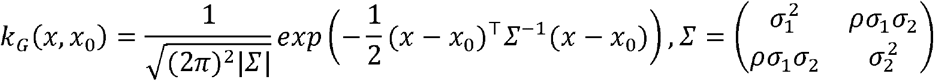

|Σ| is the determinant of the covariance matrix. We set *σ*_*1*_*=σ*_*2*_=4 and *ρ* = 1 by default.

### Dropout simulation experiments

To simulate dropout events, we first randomly selected elements in the gene expression matrix and set them to zero. The proportion of elements selected to all was defined as the dropout rate, and we set this rate in the range {0.1, 0.2, 0.3…, 0.9}. We then constructed TIST-net. To fill in missing data, we used the average expression of adjacent non-zero vertices in TIST-net. Specifically, with the increase in the dropout rate, the neighbour vertices of a missing vertex may also be lost. Therefore, we looped the fill operation above to ensure that all missed vertices are filled. Experiments show that the number of loops is positively related to the dropout rate. We calculated KL divergence between two gene expression vectors before and after the dropout operation and defined 1-KL divergence as the dropout recovery score. A higher recovery score indicates better filling results.

### Evaluation of SDE genes

We detected SDE genes for each spatial cluster by TIST. Based on the good SC identification results of TIST, we predominantly detected those genes with spatial expression patterns that were displayed in certain SCs by means of the *FindAllCluster* function in *Seurat*. We selected the top 100 genes with the largest log_2_ fold change between the mean expression value in the target SC and others using the Wilcoxon rank-sum test to test differences. We utilized the significance level defined by SPARK to rank adjusted p values for SDE genes and selected the top 100 genes for further analysis.

We reward genes with high total expression levels in adjacent spaces as more biologically meaningful and define *G*_*es*_ to quantify the spatial enrichment effect as:

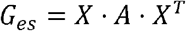

Here, *X* represents the expression vector of spots, and *A* is the adjacent matrix of KNN (the k nearest neighbours, selected according to the spatial location of the spot). This gene enrichment score is normalized to 0-1 as follows:

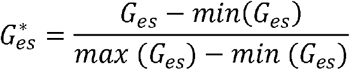

### Quantitative indicators for evaluating clustering effects

To quantitatively assess the effects of different clustering methods, we designed both supervised and unsupervised indices. With regard to supervised classification accuracy *A*_*s*_, we define:

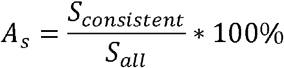

Here, *S*_*consistent*_ is the number of spots that belong to SCs identified consistent with those in the ground truth, and *S*_*all*_ is the number of all spots in a dataset. For an annotated dataset, we first obtained labels of spots belonging to different SCs by different methods and then mapped those cluster labels to the ground truth by maximum matching and calculated the matching proportion.

The unsupervised modularity *Q* score was introduced to measure the clustering effects of networks constructed by different methods. The better clustering effects present more edges within the same clusters and fewer edges among different clusters. Here, we define:

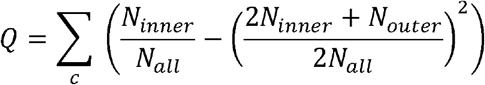

where *N*_*inner*_ and *N*_*outer*_ represent the number of edges whose vertices belong to the same clusters and to different clusters, respectively. *N*_*all*_ is the number of all edges in the network. *C* represents all clusters in the network. The value of *Q* is positively related to a better clustering effect. The range of the *Q* value is [-0.5,1). A previous study [19] showed that a *Q* range [0.3,0.7] of *Q* represented the most satisfactory clustering effect.

## Authors’ contributions

JG conceived the project. JG and QYL supervised the research. YRS and QZ conducted the research. YRS constructed TIST and finished the simulated work. QZ performed the investigation. WBG, YHW and YXM tested and improved TIST. YRS and Q.Z. wrote the original draft; WBG, YHW, YXM, HYX, QYL and JG contributed to the writing of the manuscript. All authors read and approved the final manuscript.

## Acknowledgements

This work was supported by the National Key Research and Development Program [2020YFA0712403, 2021YFF1200901]; National Natural Science Foundation of China [61922047, 81890993, 61721003, 62133006] and BNRIST Young Innovation Fund [BNR2020RC01009].

